# How do three cytosolic glutamine synthetase isozymes of wheat perform N assimilation and translocation?

**DOI:** 10.1101/733857

**Authors:** Yihao Wei, Xiaochun Wang, Zhiyong Zhang, Shuping Xiong, Yiming Zhang, Lulu Wang, Xiaodan Meng, Jie Zhang, Xinming Ma

## Abstract

To understand how the three cytosolic glutamine synthetase (GS1) isozymes of wheat (*Triticum aestivum L*., TaGS1) perform nitrogen assimilation and translocation, we studied the kinetic properties of TaGS1 isozymes, the effects of nitrogen on the expression and localization of TaGS1 isozymes with specific antibodies, and the nitrogen metabolism. The results showed TaGS1;1, the dominant TaGS1 isozyme, had a high affinity for substrates, and was widely localized in the mesophyll cells, root pericycle and root tip meristematic zone, suggesting it was the primary isozyme for N assimilation. TaGS1;2, with a high affinity for Glu, was activated by Gln, and was mainly localized in the around vascular tissues, indicating that TaGS1;2 catalyzed Gln synthesis in low Glu concentration, then the Gln returned to activate TaGS1;2, which may lead to the rapid accumulation of Gln around the vascular tissues. TaGS1;3 had low affinity for substrates but the highest V_max_ among TaGS1, was mainly localized in the root tip meristematic zone; exogenous NH_4_^+^ could promote TaGS1;3 expressing, indicating that TaGS1;3 could rapidly assimilate NH_4_^+^ to relieve NH_4_^+^ toxicity. In conclusion, TaGS1;1, TaGS1;2 and TaGS1;3 have different role in N assimilation, Gln translocation and relieving ammonium toxicity, respectively, and synergistically perform nitrogen assimilation and translocation.

**Highlight:** Three cytosolic glutamine synthase isozymes of wheat have different role and synergistically perform nitrogen assimilation and translocation.

## Introduction

Wheat (*Triticum aestivum* L.) is one of the three main cereals cultivated worldwide. Nitrogen (N) is an important limiting factor for the yield and quality of wheat, and large quantities of nitrogen fertilizers are required to attain maximal growth and productivity (Kaur *et al.*, 2015; Kichey *et al.*, 2006). To increase crop production in line with human population growth, nitrogen fertilizers are being applied excessively, leading to severe nitrogen pollution on a global scale (Kant *et al.*, 2010; Robertson and Vitousek, 2009). Therefore, there is a need to improve nitrogen use efficiency (NUE) to make agriculture more sustainable (Kant *et al.*, 2010; Thomsen *et al.*, 2014).

In order to improve crop NUE, glutamine synthetase (GS; EC 6.3.1.2) has been studied numerous times owing to its essential role in the assimilation of inorganic N (Bernard *et al.*, 2008; Fuentes *et al.*, 2001; Kichey *et al.*, 2006; Martin *et al.*, 2006; Nigro *et al.*, 2016; Tobin *et al.*, 1985). Understanding the physiological functions of GS is crucial to modulate nitrogen metabolism and to screen for germplasm with enhanced NUE (Bernard *et al.*, 2008). Plant GS is classified into two groups according to its subcellular location: the cytosolic glutamine synthetase isoform (GS1) and the chloroplastic glutamine synthetase isoform (GS2) (Goodall *et al.*, 2013; Sun *et al.*, 2015). GS2 is encoded by a single gene and plays a clear role in assimilating ammonium (NH_4_^+^) derived from photorespiration and nitrate (NO_3_^-^) reduction (Wallsgrove *et al.*, 1987), while GS1 is encoded by a multigene family and plays nonredundant and complex roles related to N assimilation and recycling (Bernard and Habash, 2009).

In Arabidopsis, GS1 is encoded by five individual isogenes with distinct affinities for NH_4_^+^ and glutamate and tissue localization as well as distinct physiological functions (Guan *et al.*, 2016; Guan *et al.*, 2015; Ishiyama *et al.*, 2004b; Konishi *et al.*, 2017; Lothier *et al.*, 2011; Moison *et al.*, 2018). Phylogenetically, the nucleotide and amino acid sequences of these isoforms do not cluster with GS1 sequences from cereals (Thomsen *et al.*, 2014). The function of Arabidopsis and cereal GS1 isogenes can thus not be compared directly, highlighting the importance of studying crop species to improve crop NUE.

Expression analyses and knockout studies of individual GS1 isogenes have demonstrated that they have specific spatial distribution and play essential roles in plant development and yield structure in rice (*Oryza sativa*), and maize (*Zea mays*) (Thomsen *et al.*, 2014). Rice has three isogenes for *GS1* (*OsGS1;1-3*). Knockout of *OsGS1;1*, which localizes to vascular tissues of mature leaves, showed severe retardation in growth rate and grain filling (Kusano *et al.*, 2011; Tabuchi *et al.*, 2005). Knockout of *OsGS1;2*, which localizes to surface cells of roots in an NH_4_^+^-dependent manner, showed a marked decrease in contents of Gln and asparagine (Asn), but increase in NH_4_^+^ content in the root and xylem sap, indicating that OsGS1;2 is important in the primary assimilation of NH_4_^+^ taken up by rice roots (Funayama *et al.*, 2013; Ishiyama *et al.*, 2004a). Real time PCR indicated that *OsGS1;3* is mainly expressed in the spikelet, indicating that it is probably important in grain ripening and/or germination (Yamaya and Kusano, 2014).

Maize has five isogenes for *GS1* (*ZmGln1;1-5*), but only *ZmGln1-3* and *ZmGln1-4* have been study so far. *ZmGln1-3* in the mesophyll cells is constitutively expressed until a very late stage of leaf development, indicating a role in the synthesis of Gln following NO_3_^-^ reduction until plant maturity (Hirel *et al.*, 2005; Martin *et al.*, 2006). *ZmGln1-4* in the bundle sheath cells was up-regulated in older leaves, indicating a role in the reassimilation of NH_4_^+^ released during protein degradation in senescing leaves (Martin *et al.*, 2005; Martin *et al.*, 2006). Furthermore, knockout of *ZmGln1-3* and *ZmGln1-3* results in reduced kernel number and kernel size, respectively (Cañas *et al.*, 2010; Martin *et al.*, 2006).

However, it is hard to study the function of TaGS1 isozymes in the allohexaploid wheat using gene knockout technology. Therefore, both the precise functions of individual TaGS1 isozymes and how they perform nitrogen assimilation and translocation are not clear. This makes it difficult to achieve goal of improving wheat NUE. On the bases of phylogenetic studies and mapping data in wheat, ten GS cDNA sequences are classified into four subfamilies denominate GS1 (a, b, and c), GS2 (a, b, and c), GSr (1 and 2), and GSe (1 and 2) (Bernard *et al.*, 2008; Thomsen *et al.*, 2014). We re-named GS1, GSr, and GSe genes as TaGS1;1, TaGS1;2, and TaGS1;3, respectively, according to phylogenetic tree analysis (Thomsen *et al.*, 2014). TaGS1;1 transcript is present in the perifascicular sheath cells, increase from anthesis, and can be upregulated in response to a reduction in N supply. Contrastingly, TaGS1;2 transcripts are confined to the vascular cells, remain at steady levels until a late stage of development, and are downregulated under N starvation (Bernard *et al.*, 2008; Caputo *et al.*, 2009). During leaf senescence, TaGS1;1 and TaGS1;2 are the predominant isoforms, suggesting major roles in assimilating ammonia during the critical phases of remobilization of nitrogen to the grain (Bernard *et al.*, 2008). The transcription level of TaGS1;3 is very low compared to other TaGS1 genes and its functions are still not known (Bernard *et al.*, 2008; Caputo *et al.*, 2009).

Since TaGS1 genes are highly homologous and their gene products were indistinguishable at the protein levels by GS antibodies, previous studies about individual TaGS1 isozyme only focus on the transcription level (Bernard *et al.*, 2008; Caputo *et al.*, 2009; Goodall *et al.*, 2013; Zhang *et al.*, 2017). However, as an enzyme, GS catalytic activity requires a process going from DNA, mRNA, and protein to subunit assembly into holoenzymes. The regulation of each step of these processes will affect the GS activity (Thomsen *et al.*, 2014). Therefore, studying the localization and expression pattern of individual GS isozyme at the protein level will be more conducive to understanding its function. Studies on the kinetic properties of individual GS isozymes in Arabidopsis and rice showed that the affinity to substrates and the ability to synthesize Gln significantly differed among GS isozymes (Ishiyama *et al.*, 2004a; Ishiyama *et al.*, 2004b). However, the kinetic properties of the individual GS1 isoenzymes have not been well characterized in wheat. In order to better understand the precise functions of individual TaGS1 isozymes and how they perform nitrogen assimilation and translocation, we studied the kinetic properties of TaGS1 isozymes, the effects of nitrogen on the expression and localization of TaGS1 isozymes with specific antibodies, and the nitrogen metabolism. Based on these new data, we discovered that three TaGS1 isozymes have different role and synergistically perform nitrogen assimilation and translocation. The results of this study are expected to provide directions for improving nitrogen use efficiency of wheat, which would in turn, abate N pollution arising from the use of excess N fertilizers.

## Materials and methods

### Expression of recombinant wheat GS protein in *E. coli*

The CDS (Coding Sequence) region of *TaGS1;1, TaGS1;2, TaGS1;3*, and *TaGS2* were obtained from wheat cultivar Yumai 49, and they were respectively cloned into the pET21a vector in our previous study (Gu *et al.*, 2018). The recombinant wheat GS protein was induced in accordance with a method described by Gu *et al.* (2018). After induction, cells were harvested by centrifugation at 5000 g for 10 min at 4 °C. The pellet was suspended in breaking buffer (10 mmol/L Tris, 10 mmol/L MgCl_2_, 0.05 % Triton X-100, 100 μg/mL PMSF, pH 7.5) and sonicated using an ultrasonic homogenizer JY92-2D (Ningbo Scientz Biotechnology Co. Ltd., Ningbo, China). The lysate was centrifuged at 12000 g for 15 min at 4 °C and the supernatants were collected and used for kinetic measurements. Supernatant was kept on ice until used.

GS abundance in the supernatant was detected using Western-blot. GS polypeptides were detected using polyclonal antibodies raised against both wheat GS1 and GS2. The relative content of GS was estimated by grey scanning using Image Lab analyzer software (Version 5.1, Bio-Rad Laboratories, CA, USA).

### In vitro assay of individual recombinant wheat GS isozymes activity

Determination of GS enzyme activity was based on an in vitro modified synthetase reaction, where the amount of produced γ-glutamyl monohydroxamate (GMH) is detectable by a stop reaction (Ma *et al.*, 2005; Németh *et al.*, 2018).

The crude extract of wheat GS protein recombinant *E. coli* was added into 800 μL Reagent buffer. The reaction mixture was incubated at 25 °C for 15 min, terminated by adding 800 μL stop solution (123 mM FeCl_3_, 49 mM trichloroacetic acid, and 217 mM HCl) after centrifuging at 12000 g for 5 min, and the absorbance of supernatants at 540 nm was determined. Reagent buffer always contained 40 mM magnesium sulfate and basically 100 mM imidazole, 50 mM ATP, 40 mM hydroxylamine, and 50 mM Na-glutamate but concentrations varied depending on the actual kinetic assay (i.e., Na-glutamate: 0–120 mM; glutamine: 0–60 mM; hydroxylamine: 0–80 mM).

### Design and preparation of antibodies against individual wheat GS isozymes

DNA Star was used to compare the amino acid sequences of TaGS1;1, TaGS1;2, TaGS1;3, and TaGS2 (Fig. S1). The hydrophilic and surface accessibility and antigenicity of these polypeptide sequences with low homology were analyzed with Protean (Table S1). The polypeptide sequences with strong antigenic, hydrophilic, and surface accessibility were selected as individual wheat GS isozymes antigenic polypeptides, i.e., TaGS1;1: KDGGFKVIVDAVEKLKLKHKE; TaGS1;2: EAGGYEVIKTAIEKLGKRHAQ; TaGS1;3: LSKAGLSNGK; TaGS2: TLEAEALAAKKLALKV. These antigenic polypeptides were synthesized and polyclonal antibodies against individual wheat GS isozymes were prepared by Zoonbio (Zoonbio Biotechnology Co., Ltd., Nanjing, China). To prepare polyclonal antibodies against wheat GS protein, the purified recombinant protein antigen of TaGS1;1 and TaGS2 was used.

### Plants growth conditions and experimental design

For hydroponic treatments, uniform seeds were selected, surface sterilized with 75 % (v/v) ethanol for 1 min, rinsed with distilled water, and then germinated in culture dishes covered with wet sterilized filter paper until the seed root length was about 1 cm. The uniform seedlings were transplanted to opaque containers and cultivated in distilled water. The hydroponic culture was carried out in a growth chamber with the following conditions: 22 °C ± 2 °C, 50 % to 70 % relative humidity, a photon fluence rate of 300 μmol photons m^−2^ s^−1^, and a 16h light period. After three days, the seedlings were separated and grown on a modified Hoagland nutrient solution (Table S2), with NH_4_^+^ or NO_3_^-^ as the sole N source at concentrations of 0, 0.2, 2, 5, 10, and 20 mM. Each container contains 10 plants with 0.5 L nutrient, which was replaced every 3d. After 12 days, the shoots and roots were harvested individually and immediately frozen in liquid nitrogen, then stored at −80 °C for further experiments. In parallel, leaves and roots were selected, and immediately immersed in fixative for the immunolocalization studies.

### RNA Isolation and quantitative real-time PCR

Total RNA was extracted from the plant tissue using TRIzol Reagent (Thermo Scientific). The cDNA was synthesized using the Hiscript 1st Strand cDNA Synthesis Kit (Vazyme). Quantitative real-time PCR (qPCR) was performed on Step One Real-Time PCR System (Life Technologies Corporation, CA, USA), and AceQ qPCR SYBR Green Master Mix (Vazyme) for the assay. All the primers (Sangon) used are shown in Table S3. The relative expression levels of the genes were calculated using the TaATPases (Ta54227) and TaTEF (Ta53964) gene (Paolacci *et al.*, 2009) as internal control.

### GS Activity Assay and Western blotting

The total GS activity was measured in accordance with a method described by Ma *et al.* (2005). Soluble protein content was determined by the Coomassie blue dye-binding method using bovine serum albumin as a standard. Native-polyacrylamide gel electrophoresis (PAGE) and in-gel GS activity staining were performed as previously described (Zhang *et al.*, 2017). Western blotting was performed in accordance with a method described by Wei *et al.* (2018). The dilution ratio of antibody applied to the membrane is indicated in the figure legends.

### Metabolite Analysis

Amino acid, ammonium, and nitrate were determined according to Wei *et al.* (2018). Total N content determined using SEAL AutoAnalyzer 3 continuous flow analytical system (Bran + Luebbe, Hamburg, Germany), in accordance with the manufacturer’s instructions. Soluble sugar was determined using the anthrone colorimetric method (Tang, 1999).

The amino acid components were analyzed with thin layer chromatography (TLC). Amino acids were identified with phenol-water (3:1) as a developing solvent in silica gel G plate and using 0.5 % ninhydrin n-butyl alcohol solution as the visualization reagent. Total free amino acid extracted from the shoot or root tissues (1.5 μg) were loaded onto each lane of the TLC.

### Immunolocalization using indirect immunofluorescence analysis

These tissues of leaf, root and root tip were fixed in FAA fixative at least 24h. Embedded in paraffin, section and immunofluorescence were prepared by Servicebio (Wuhan Servicebio technology Co., Ltd., Hubei, China). Anti-TaGS1;1, anti-TaGS1;2, anti-TaGS1;3, and anti-TaGS2 antibodies diluted 1:200, 1:200, 1:500 and 1:50, respectively, in blocking solution.

### Statistics

One-way analysis of variance with a Duncan post hoc test was performed using SPSS version 13.0 (IBM, Chicago, IL, USA).

### Accession numbers

The locus numbers for wheat GS cDNA are as follows: *TaGS1;1* (DQ124209), *TaGS1;2* (AY491968), *TaGS1;3* (AY491970), *TaGS2* (DQ124212).

## Results

### The individual wheat GS isozymes are distinguishable at the protein level with TaGS specific antibody

The individual recombinant wheat GS protein loaded in the gel was adjusted to a uniform level using polyclonal antibodies raised against wheat GS (Fig. 1A). Then, the specificity of antibodies against individual wheat GS isozymes was tested. The results showed that anti-TaGS1;1, anti-TaGS1;2, anti-TaGS1;3, and anti-TaGS2 antibodies were monospecific to TaGS1;1, TaGS1;2, TaGS1;3, and TaGS2 polypeptides, respectively, with an antibody dilution ratio of 1:30000, 1:30000, 1:10000, and 1:10000, respectively (Fig. 1A).

**Fig. 1.**
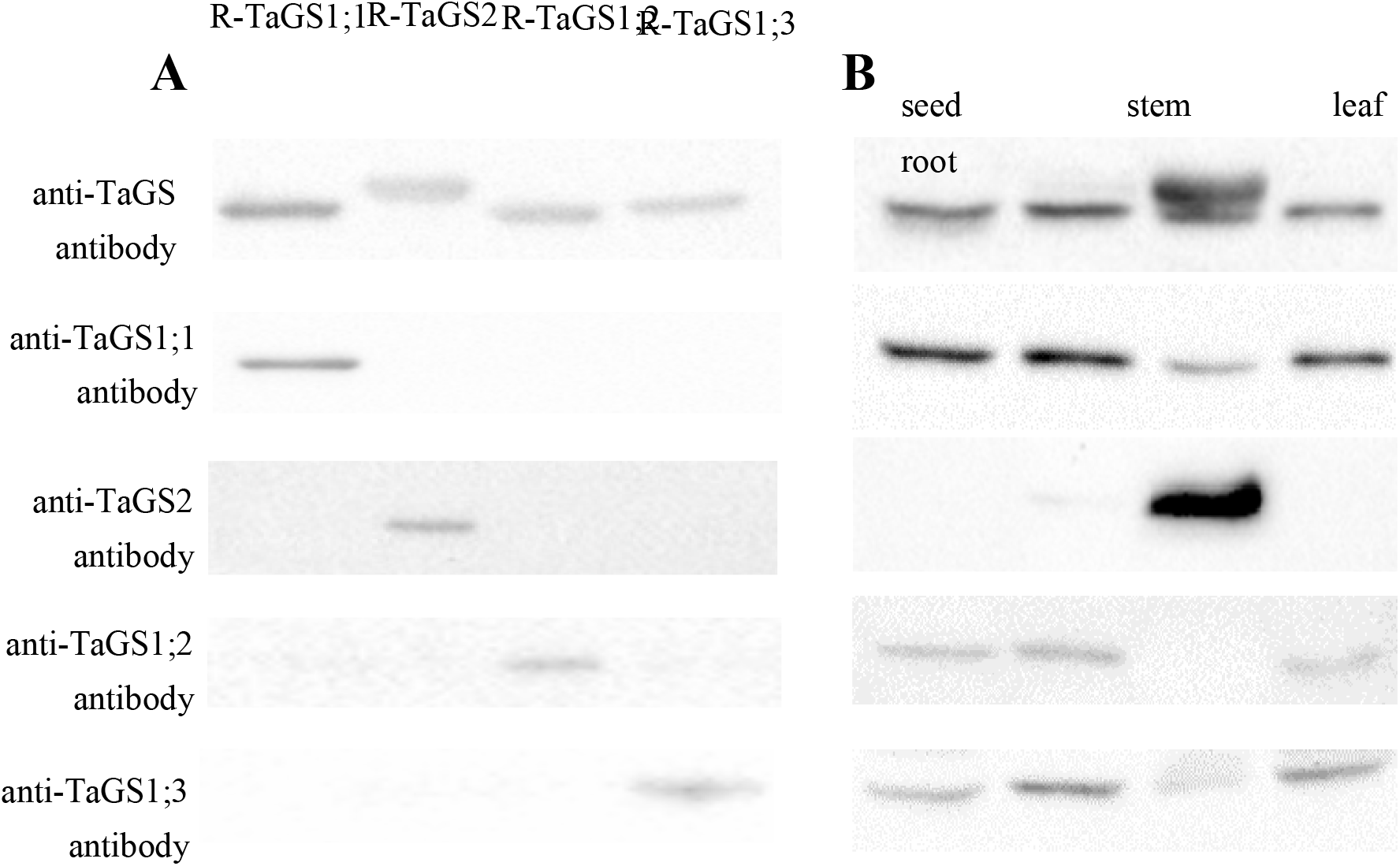
Cross-reactivity of anti-GS antibodies to the individual recombinant wheat GS proteins (A) and the wheat GS proteins in different tissues (B). The dilution ratio of the anti-TaGS, anti-TaGS1;1, anti-TaGS2 anti-TaGS1;2 and anti-TaGS1;3 antibody, is 1:5000, 1:30000, 1:10000, 1:30000 and 1:10000, respectively.

The anti-TaGS2 antibody was monospecific to TaGS2 polypeptide for only one band detected at about 42 kDa when wheat stem and leaf extract were analyzed by immunoblotting after SDS-PAGE; the antibodies of individual TaGS1 did not cross-react against TaGS2 polypeptide, as only one band at about 39 kDa was detected when wheat stem and leaf extract were analyzed by immunoblotting (Fig. 1B).

### Kinetic properties of recombinant TaGS isoforms

Kinetics of GS activities of recombinant TaGS1;1, TaGS1;2, TaGS1;3, and TaGS2 were plotted against the concentration of glutamate (Glu), hydroxylamine, and Gln in the reaction mixture (Fig. 2). TaGS1;2 activity was significantly inhibited when Glu was supplied at the concentrations higher than 6 mM (Fig. 2A), and was very weak at the different concentrations of hydroxylamine when Glu was supplied at the 50 mM (Fig. 2B). However, TaGS1;2 activity was not inhibited when Glu was supplied at concentrations lower than 5 mM (Fig. 2D) and was significantly increased at the different concentrations of hydroxylamine when Glu was supplied at the 5 mM (Fig. 2E). With increasing Gln concentrations, the activities of TaGS1;3 and TaGS2 remained stable. Contrastingly, in the reaction mixture with 60 mM Gln, the activity of TaGS1;1 and TaGS1;2 was increased to about 2 and 6 times the activity in the reaction mixture without Gln, respectively (Fig. 2C).

**Fig. 2.**
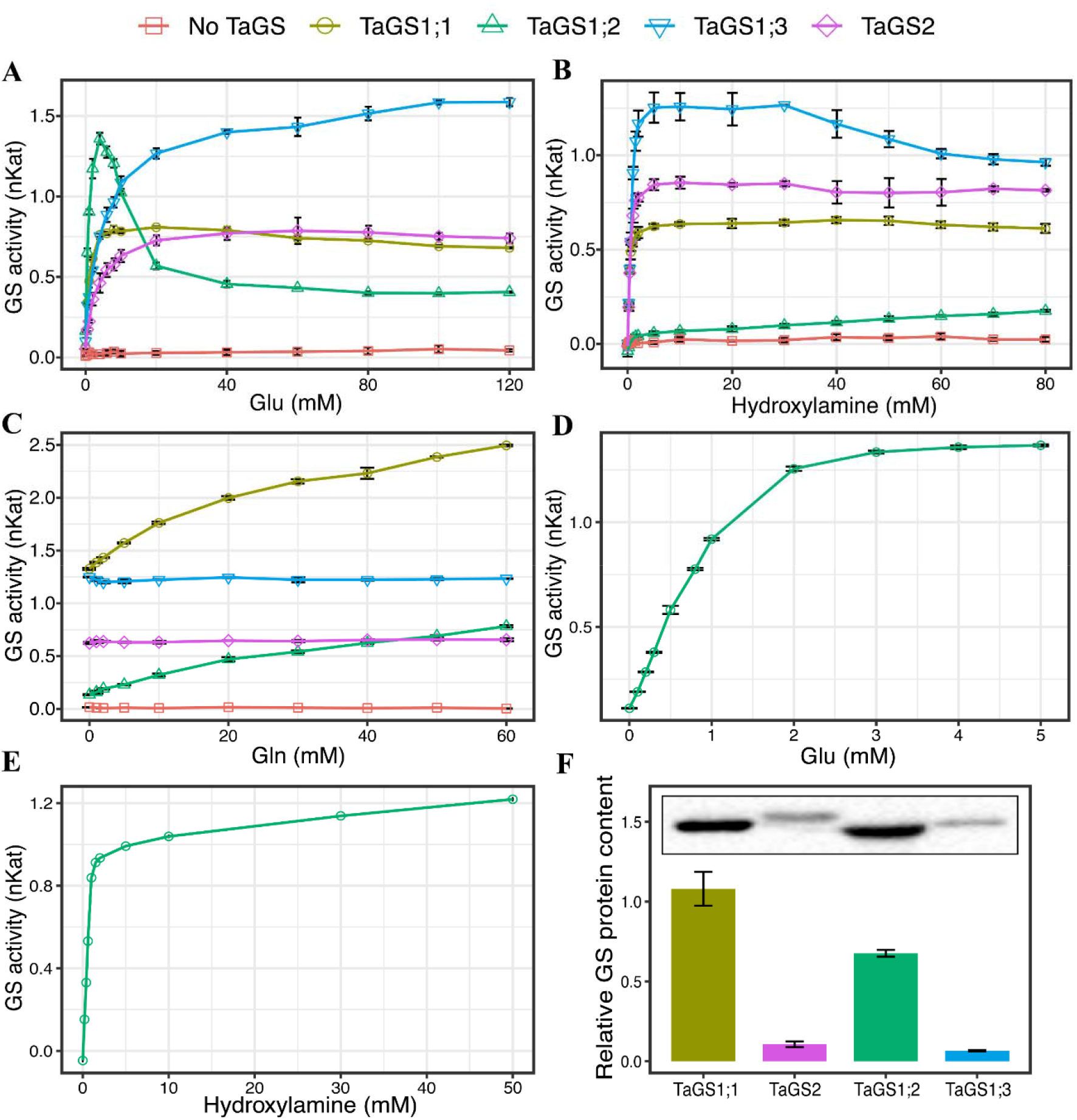
The activity of individual recombinant wheat GS isozymes in relation to additional Glu (A), hydroxylamine (B) and Gln (C). The TaGS1;2 activity was measured when Glu was supplied at the concentrations 0-5 mM (D). The TaGS1;2 activity was measured at the different concentrations of hydroxylamine when Glu was supplied at the 5mM (E). The volume of individual recombinant wheat GS isozymes crude extract, with 200 μL of TaGS1;1, 150 μL of TaGS1;2, 450 μL of TaGS1;3 and 300 μL TaGS2 were used for the GS enzyme assays. The relative GS protein content of recombinant wheat GS isozymes crude extract (F). Upper panel showed the TaGS immunoblot and lower panel showed the quantified intensity of the TaGS Western bands present in upper panel. Data represent means ± SE of at least three replicates.

The specific activities plotted against the substrate concentrations showed saturation kinetics, which followed the Michaelis-Menten equations, and the kinetic constants were calculated (Table 1). The four TaGS isoenzymes can be classified into different groups by the affinities to substrates. TaGS1;1 (K_m_ = 0.65 ± 0.01 mM) and TaGS1;2 (K_m_ = 0.87 ± 0.01 mM) can be classified as isoenzymes with high affinity to Glu, while TaGS1;3 and TaGS2 exhibited a low affinity to Glu (K_m_ values; 4.13 ± 0.35 mM and 2.43 ± 0.27 mM, respectively). As for hydroxylamine, TaGS1;1 (K_m_= 0.26 ± 0.02 mM) and TaGS2 (K_m_= 0.36 ± 0.04 mM) showed high substrate affinity than TaGS1;2 (K_m_= 0.66 ± 0 mM) and TaGS1;3 (K_m_= 0.64 ± 0.04 mM).

**Table 1.** The kinetic properties of the wheat GS isoenzymes

| | $K_m^*$ (mM) | | $V_{max}^{\#}$ (nKat/1 unit protein) | |
| --- | --- | --- | --- | --- |
|  | Glu | Hydroxylamine | Glu | Hydroxylamine |
| TaGS1;1 | 0.65±0.01 d | 0.26±0.02 c | 0.13±0.001 d | 0.1±0.002 d |
| TaGS1;2 | 0.87±0.01 c | 0.66±0.003 a | 0.5±0.002 c | 0.36±0.003 c |
| TaGS1;3 | 4.13±0.35 a | 0.64±0.04 a | 1.63±0.06 a | 1.36±0.07 a |
| TaGS2 | 2.43±0.27 b | 0.36±0.04 b | 0.79±0.03 b | 0.85±0.03 b |

Note: <sup>\*</sup> For TaGS1;1, TaGS1;2, TaGS1;3 and TaGS2, the concentration of Glu used for curve fitting was 0-40mM, 0-5 mM, 0-120 mM and 0-120 mM, respectively; the concentration of hydroxylamine used for curve fitting was 0-80 mM, 0-50 mM, 0-30 mM and 0-80 mM, respectively. The volume of individual recombinant wheat GS isozymes crude extract, with 200 μL of TaGS1;1, 150 μL of TaGS1;2, 450 μL of TaGS1;3 and 300 μL TaGS2 were used for the GS enzyme assays.
<sup>#</sup> One kat of enzyme activity was defined as 1 mol GMH synthesized per second at 25°C.
Data are means of three independent biological replicates ± SD. The different letters above each sample indicate statistically significant differences where $P < 0.05$ according to one-way ANOVA Duncan post-hoc test.

The relative GS protein content of recombinant wheat GS isozymes crude extract was determined by analyzing the Western-blot results using Image Lab (Fig. 2F), the results showed a TaGS1;1: TaGS2: TaGS1;2: TaGS1;3 ratio of 1: 0.1: 0.63: 0.06. Based on the relative content of different TaGS isozymes, we calculated the V_max_ of each TaGS isozymes. The V_max_ of TaGS1;3 was the highest, about 10-fold, 3-fold, and 2-fold of TaGS1;1, TaGS1;2, and TaGS2, respectively (Table 1).

### Effects of nitrogen nutrition on individual TaGS gene expression

The responses of GS genes to N nutrition are crucial to understand their role in N metabolism. The relative abundances of the mRNA and subunit of individual GS isozymes in wheat seedling were determined by real time PCR and Western blot analyses.

#### The expression pattern of individual TaGS at the transcript level

The TaGS1;1 transcript was the highest among all TaGS1 genes (Fig. 3A), suggesting that TaGS1;1 was the dominant TaGS1 isoform in the shoot and root. And TaGS1;1 transcript was higher under 0 - 0.2 mM N supply than under 2 - 20 mM N supply (Fig. 3A).

**Fig. 3.**
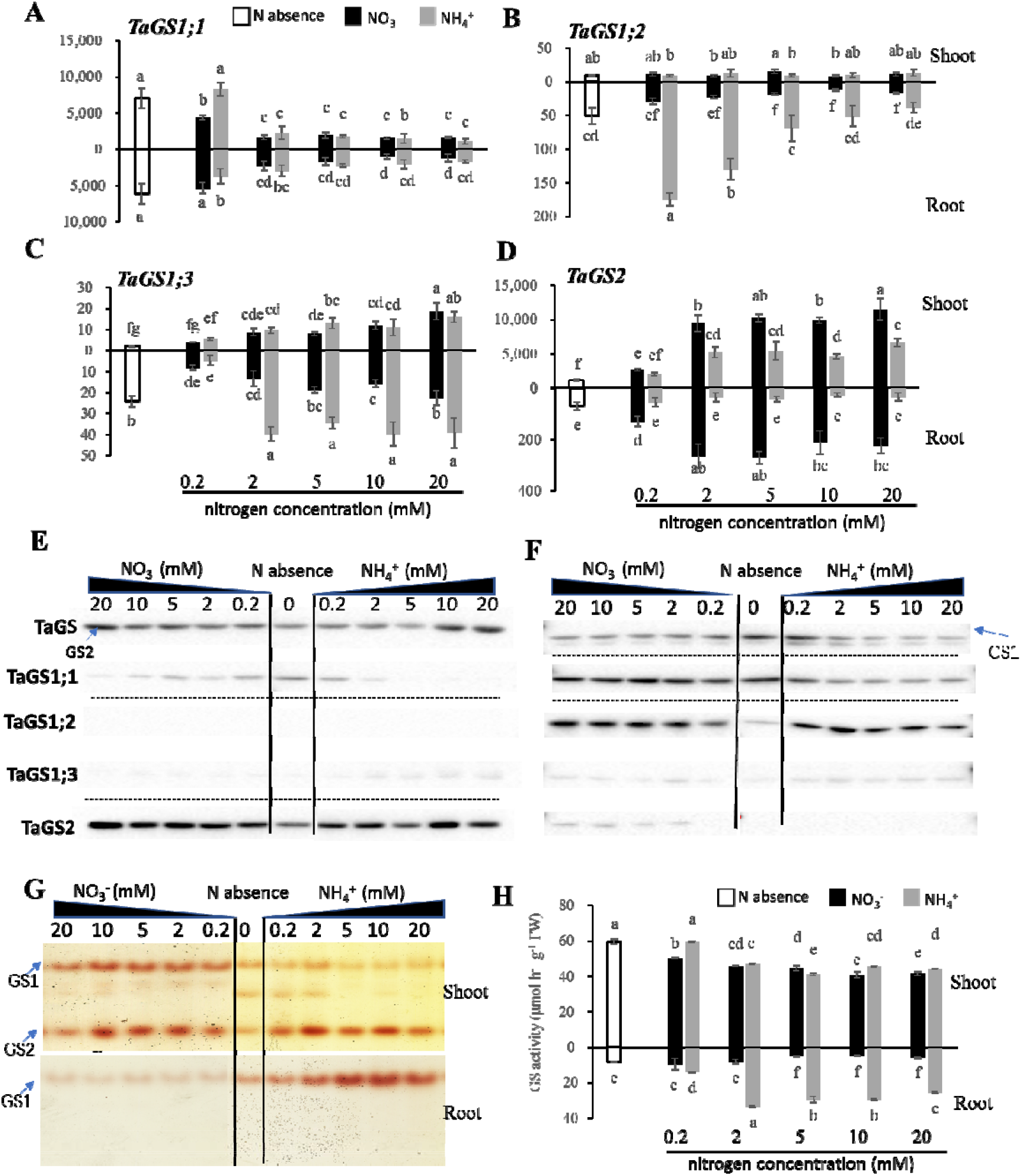
TaGS gene expression and GS activity in response to different nitrogen regimes in the shoot and root tissue. Quantitative RT–PCR analysis of TaGS1;1 (A), TaGS1;2 (B), TaGS1;3 (C) and TaGS2 (D) gene expression, horizontal axes present the N absence and the millimolar concentration of NO_3_^-^ and NH_4_^+^ treatments, vertical axes present the mean relative expression of each isoform normalized to the reference genes *TaATPase* and *TaTEF.* Western-blot analysis of TaGS, TaGS1;1, TaGS1;2, TaGS1;3 and TaGS2 protein contents in shoot (E) and root (F), 15 μg of soluble proteins extracted from the tissues were loaded onto each lane and detected by the anti-TaGS, anti-TaGS1;1, anti-TaGS1;2, anti-TaGS1;3 and anti-TaGS2 antibody, respectively and the dilution ratio of antibody applied to the membrane is the dilution ratio of antibody is 1:5000, 1:30000, 1:30000, 1:10000 and 1:10000, respectively. Native electrophoresis and in-gel GS activity staining (G) showing the GS holoenzymes in shoot and root; The total GS activity (H) in shoot and root tissue under different N regimes. Data are means of three independent biological replicates ± SD. The different letters above each sample indicate statistically significant differences where *P* < 0.05 according to one-way ANOVA Duncan post-hoc test.

The TaGS1;2 transcript in the shoot was much lower than that in root, and was not affected by N treatments (Fig. 3B). In the root, TaGS1;2 transcript was the highest under 0.2 mM NH_4_^+^ supply, and decreased with increasing NH_4_^+^ supply (Fig. 3B). However, the TaGS1;2 transcript in root did not significantly change with increasing NO_3_^-^ supply, and was significantly lower than under NH_4_^+^ supply (Fig. 3B).

The TaGS1;3 transcript increased with increasing N supply in the shoot, but was very lower than that in the root (Fig. 3C). In roots, with increasing NO_3_^-^ supply, the TaGS1;3 transcript increased, but it was significantly lower than under NH_4_^+^ supply (Fig. 3C). And the TaGS1;3 transcript in root was higher under 2 - 20mM NH_4_^+^ than under 0.2 mM NH_4_^+^ supply (Fig. 3C), indicating that high NH_4_^+^ concentration can induce the expression of TaGS1;3 in roots.

The TaGS2 transcript was much higher in the shoot than that in the root (Fig. 3D). In the shoot, TaGS2 transcript was higher under 2 - 20 mM N supply than under 0 - 0.2 mM (Fig. 3D). When under 2 - 20 mM N supply, the TaGS2 transcript was significantly higher under NO_3_^-^ than under NH_4_^+^ supply. Interestingly, when under 2 - 20 mM NO_3_^-^ supply, TaGS2 transcript in the root was significantly higher than without N or under NH_4_^+^ supply (Fig. 3D).

#### The expression pattern of individual TaGS subunit

Western blots analysis using polyclonal antibodies raised against GS of wheat, showed that GS2 was the predominant isoform in the shoot and GS1 was the predominant isoform in the root (Fig. 3E, F). In the shoot, TaGS2 subunit level increased with increasing N supply (Fig. 3E). In the root, TaGS1 subunit decreased with increasing N supply (Fig. 3F). Furthermore, TaGS2 subunit could be detected in root only when NO_3_^-^ concentration was greater than 0.2 mM, indicating that TaGS2 in the root was specifically induced by high concentration NO_3_^-^ (Fig. 3F).

In the shoot, TaGS1;1 subunit abundance decreased with increasing N supply, becoming difficult to detect when NH_4_^+^ concentration was greater than 2 mM (Fig. 3E). In the roots, TaGS1;1 subunit decreased with increasing NH_4_^+^ supply but it did not significantly change with increasing NO_3_^-^ supply (Fig. 3F). Less TaGS1;2 subunit was detected in the shoot while much TaGS1;2 were detected in the root. Furthermore, TaGS1;2 subunit increased with increasing NO_3_^-^ supply in the root, but it first increased and then decreased with increasing NH_4_^+^ supply (Fig. 3E, F). The TaGS1;3 subunit was very low in the shoot and root, and it was higher under NH_4_^+^ than under NO_3_^-^ supply (Fig. 3E, F).

### Effects of nitrogen nutrition on GS isozymes activity and total GS activity

In previous studies, the cytosolic GS1 holoenzyme was ~490 kDa, and the chloroplastic GS2 holoenzyme was ~240 kDa (Wang *et al.*, 2015). Therefore, the isoforms showed different mobilities in gels (GS2 > GS1). In the shoot, the activity of both GS1 and GS2 isozymes could be detected, but only GS1 isozymes activity could be detected in the root (Fig. 3G). GS1 activity in the shoot was significantly higher under NO_3_^-^ than under NH_4_^+^ supply. However, GS1 activity in the root was significantly lower under NO_3_^-^ than under NH_4_^+^ supply (Fig. 3G). Moreover, the GS1 activity in the root reached its peak at 10 mM NH_4_^+^ (Fig. 3G).

The total GS activity in the shoot was significantly higher than that in the root, and the total GS activity in the root was significantly higher under NH_4_^+^ than under NO_3_^-^ supply, and increased significantly by high concentration NH_4_^+^ (Fig. 3H).

### Effects of nitrogen nutrition on C/N metabolite status

Without N supply, the shoot growth was significantly inhibited, while the root growth was significantly promoted (Table S4). The free NH_4_^+^ producing by its own metabolic process, was significantly higher in the root than in the shoot (Fig. 4A). Soluble sugar, the main product of photosynthesis, also accumulated in the roots (Fig. 4B). In the root and shoot, the free amino acid (Fig. 4C), soluble protein (Fig. 4D), and total nitrogen content (Table S4) were lower than those under nitrogen sufficiency, showing that nitrogen assimilation was inhibited under nitrogen deficiency.

**Fig. 4.**
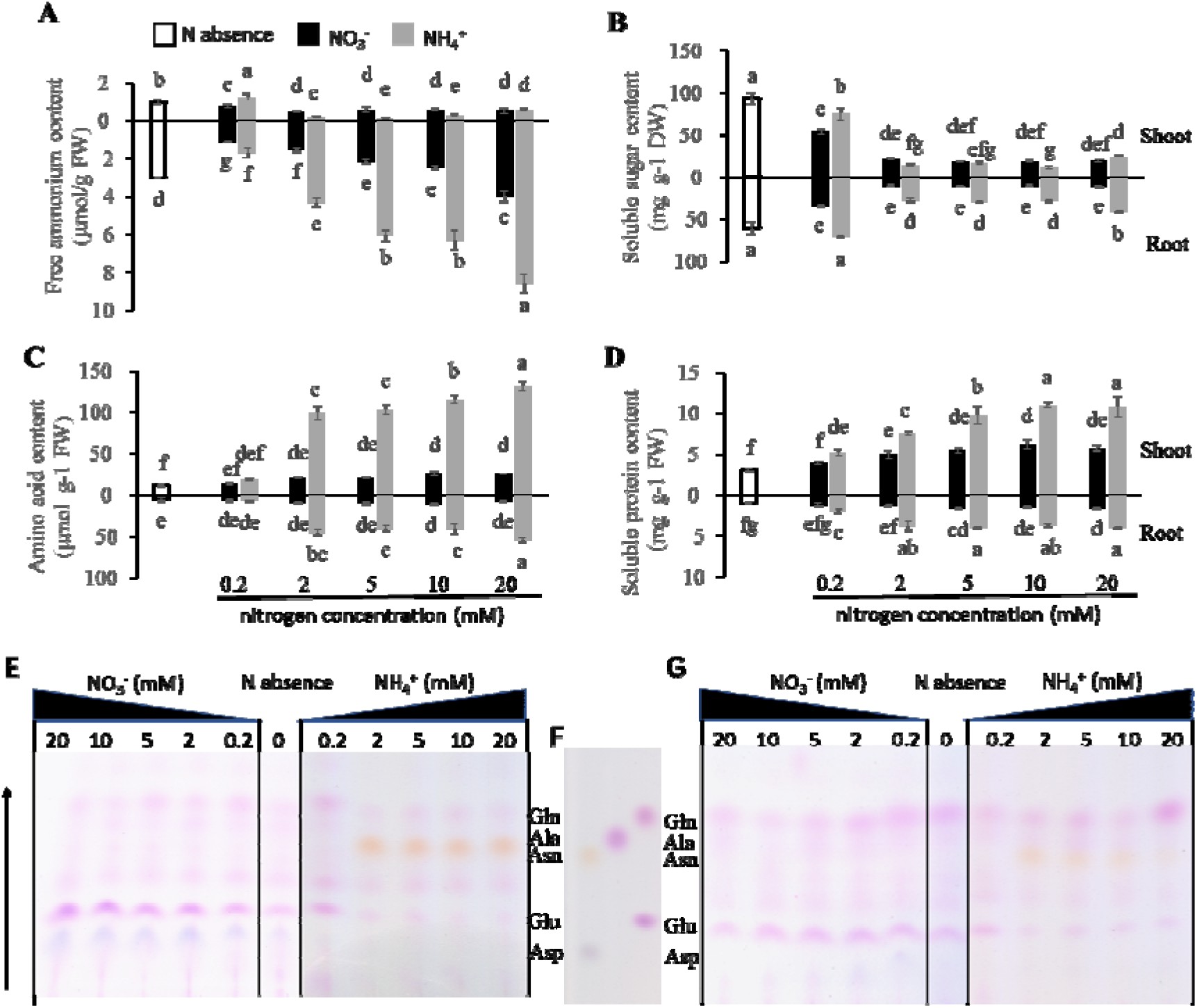
Carbon and nitrogen metabolite levels in response to different nitrogen regimes in the shoot and root tissue. The ammonium (A), soluble sugar (B), free amino acid (C) and soluble protein content (D) were determined. Individual amino acid components in response to different nitrogen regimes in the shoot and root tissue. The amino acid components were analyzed with thin layer chromatography (TLC) and ninhydrin colouring showing the amino acid components in shoot (E) and root (G). F showed the separation results of five amino acids (Gln, Ala, Asn, Glu and Asp) under the same conditions. 1.5 μg of free amino acid extracted from the shoot and root tissues were loaded onto each lane of the TLC. Data are means of three independent biological replicates ± SD. The different letters above each sample indicate statistically significant differences where P < 0.05 according to one-way ANOVA Duncan post-hoc test.

Under 0.2 mM NO_3_^-^ supply, NO_3_^-^ was preferentially accumulated in the shoot (Fig. S2). With increasing NO_3_^-^ concentration, the content of NO_3_^-^ in the shoot and root and the content of free NH_4_^+^ in the root gradually increased (Fig. S2 and Fig. 4A), but the content of the organic nitrogen (free amino acid, and soluble protein) did not show an increasing trend (Fig. 4C, D), which indicated that nitrogen was mainly stored as inorganic nitrogen with increasing NO_3_^-^ supply.

Under NH_4_^+^ supply, the content of free NH_4_^+^ in the root was significantly higher than in the shoot and increased with increasing NH_4_^+^ supply (Fig. 4A), but the content of organic nitrogen (free amino acid, and soluble protein) in the shoot was significantly higher than in the root and increased with increasing NH_4_^+^ supply (Fig. 4C, D), suggesting that NH_4_^+^ was mainly stored in the root while organic nitrogen was mainly stored in the shoot with increasing NH_4_^+^ supply.

Under 2 - 20 mM N supply, the content of soluble sugar in the root was higher than that in the shoot under NH_4_^+^ supply, but it was lower than in the shoot under NO_3_^-^ supply (Fig. 4D), indicating that the root needs more carbohydrates to assimilate NH_4_^+^ under NH_4_^+^ supply. From 2 to 20 mM nitrogen, the amino acids content in plant under NH_4_^+^ supply was about 2–3 times that under NO_3_^-^ supply and the amino acids contents in the shoot gradually increased with increasing NH_4_^+^ supply (Fig. 4C). Moreover, the soluble protein content in plant under NH_4_^+^ supply was significantly higher than under NO_3_^-^ supply (Fig. 4D). These results indicate that the N assimilation was enhanced in NH_4_^+^-fed wheat.

The components of free amino acid extracted from the shoot and root tissues were separated with thin layer chromatography (TLC) and stained with ninhydrin, and the main components were Gln, asparagine (Asn), Glu, and aspartate (Asp) (Fig. 4E, G). The relative proportion of Glu and Asp in the shoot and root were significantly higher under NO_3_^-^ than under NH_4_^+^ supply. However, the relative proportion of Gln and Asn in the shoot and root were significantly lower under NO_3_^-^ than under NH_4_^+^ supply (Fig. 4E, G).

### Effect of nitrogen on the tissue localization of individual TaGS

Responses of tissue localization of TaGS to N nutrition are crucial to understand the role of individual TaGS in N metabolism. Without N supply, TaGS2 and TaGS1;1 were the main isoforms and localized in the leaf mesophyll cells and TaGS1;1 was also localized in the surrounding vessels of xylem in the vein of leaf (Fig. 5A). TaGS1;2 was mainly localized in the surrounding vessels of xylem while no obvious TaGS1;3 was detected in the leaf (Fig. 5A). Only TaGS1;1 was detected in the vascular bundles in the maturation zone of roots (Fig. 6B), but abundant TaGS1;1 and TaGS1;3 were found in the meristematic zone of roots (Fig. 5C).

**Fig. 5.**
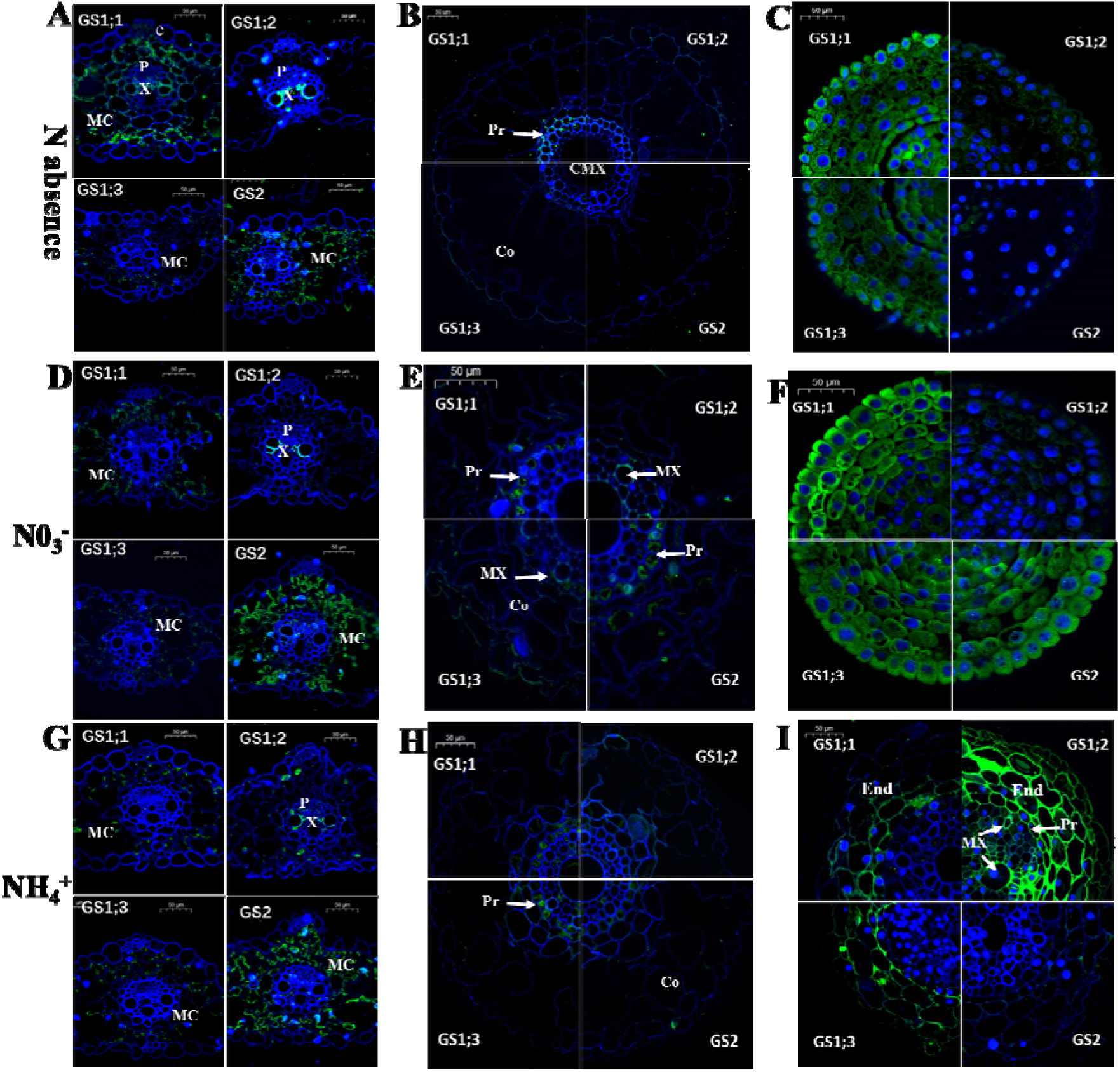
Tissue localization of individual TaGS. Immunolocalization of TaGS1;1, TaGS1;2, TaGS1;3 and TaGS2 in response to different nitrogen regimes in a transverse section of the leaf (A, D, G), the maturation zone (B, E, H) and meristematic zone (C, F, I) of root tissue. DAPI glowed blue by UV excitation wavelength 330-380 nm and emission wavelength 420 nm; FITC glowed green by excitation wavelength 465-495 nm and emission wavelength 515-555 nm. e, epidermis, MX, metaxylem; P, phloem; X, xylem; VB, vascular bundle; CMX, central metaxylem; End, endodermis; Pr, pericycle; Co, cortex; MC, mesophyll cells.

Under 5 mM NO_3_^-^ supply, the tissue localization of individual TaGS in the leaves was very similar to that without N supply, but there were more TaGS2 and less TaGS1;1 in the mesophyll cells and no TaGS1;1 was detected in the surrounding vessels of xylem (Fig. 5D). In the maturation zone of roots, TaGS1;1 and TaGS2 were localized in the pericycle cell, and TaGS1;2 and TaGS1;3 were localized in the surrounding vessels of xylem (Fig. 5E). Moreover, abundant TaGS1;1, TaGS1;3, and TaGS2 were detected in the meristematic zone of roots (Fig. 5F).

Under 5 mM NH_4_^+^ supply, the tissue localization of TaGS1;1, TaGS1;3, and TaGS2 in the leaves was the same with 5 mM NO_3_^-^ supply. TaGS1;2 was localized in the surrounding vessels of xylem and phloem companion cells in the leaves (Fig. 5G). In the maturation zone of roots, TaGS1;1 and TaGS1;3 were localized in the pericycle cells, and TaGS1;2 and TaGS2 were not detected (Fig. 5H). In the corresponding site of root tips in other treatments, the supposed root tip meristem under NH_4_^+^ treatment was full of vascular tissue (Fig. 5I). There were abundant TaGS1;2, TaGS1;3, and TaGS1;1 in the endodermis, but no TaGS2 was detected (Fig. 5I). Moreover, TaGS1;2 was detected in the surrounding vessels of xylem in the vascular bundle (Fig. 5I).

## Discussion

This study was conducted to improve our understanding of the function of glutamine synthetases, which are critical in nitrogen metabolism in wheat, to be able to provide directions of improving nitrogen use efficiency in plants. NO_3_^-^, as an important source of nitrogen absorbed by wheat roots, was reduced into NH_4_^+^ in leaves, and then was assimilated by GS2 in the chloroplast (Goodall *et al.*, 2013; Lothier *et al.*, 2011; Sivasankar and Oaks, 1996). Previous studies showed that the GS2 transcripts were in roots (Bernard *et al.*, 2008; Goodall *et al.*, 2013; Wei *et al.*, 2018), but it was first time for us to discover that TaGS2 peptide and transcripts in wheat roots were induced by NO_3_^-^ (Fig. 3D, F), and was localized in the pericycle cells (Fig. 5E) and the meristematic zone (Fig. 5F) of roots only under NO_3_^-^ supply, which indicated that TaGS2 mainly participated in assimilation of ammonia from NO_3_^-^ reduction in the root.

It is important to study the enzymatic kinetics of TaGS isozymes to illustrate their biological function. Our study showed that recombinant TaGS1 isozymes had significantly different enzymatic kinetics. TaGS1;1 is one major isoform in wheat seedling (Fig. 3A, E, F), it showed high substrate affinity for Glu and hydroxylamine but its maximum reaction rate was lowest (Table 1). Previous studies showed that TaGS1;1 transcripts are present in the perifascicular sheath cells (Bernard *et al.*, 2008), but we found that TaGS1;1 peptide was mainly localized in the leaf mesophyll cells (Fig. 5A, D, G). In the mesophyll cells, ammonium released in mitochondria during photorespiration is reassimilated in the chloroplast by GS2 (Wallsgrove *et al.*, 1987). Oliveira *et al.* (2002) found that overexpression of cytosolic GS1 in leaf mesophyll cells seems to provide an alternate route to chloroplastic GS2 for the assimilation of photorespiratory ammonium. therefore, we speculate that TaGS1;1 in mesophyll cells may participate in the reassimilation of ammonium released during photorespiration. In addition, TaGS1;1 was also localized in the pericycle and the meristematic zone of roots (Fig. 5), suggesting that TaGS1;1 was involved in assimilating inorganic nitrogen in roots and assimilating NH_4_^+^ from the root tip metabolism process. The very wide tissue distribution and high expression of TaGS1;1 indicated that it was the primary TaGS1 isozyme for N assimilation.

A recent study showed that *AtGln1;3*, located in the pericycle of root in the Arabidopsis, is involved in xylem loading of Gln (Konishi *et al.*, 2017). But how it participates in the process is unclear. In the reaction mixture with 60 mM Gln, the activity of TaGS1;2 was increased to about 6 times of that without the Gln (Fig. 2C). As GS catalytic product, Gln has no feedback inhibition effect on TaGS1;2, but significantly enhanced the catalytic activity of TaGS1;2. Furthermore, TaGS1;2 has a high affinity for Glu (Table 1), which may indicate that TaGS1;2 catalyzes rapidly the synthesis of Gln, leading to the accumulation of Gln. TaGS1;2 was mainly localized around the vascular tissues (Fig. 5G, I), suggesting that TaGS1;2 can promote the accumulation of Gln around vascular tissues. Gln is the main translocation form of plant organic nitrogen (Setién *et al.*, 2013). In wheat, Gln concentration in leaf phloem sap was dozens of times higher than in leaf tissue, where it was preferentially loaded into the vascular tissue for translocation (Duan *et al.*, 2000). Peeters and Van Laere (1994) pointed out that Gln has an amazing reverse concentration loading efficiency to vascular tissue. Therefore, it can be considered that TaGS1;2, which is mainly distributed around the vascular tissues, can rapidly accumulate Gln around the vascular tissues, thus promoting the reverse concentration loading of Gln to the vascular tissues. TaGS1;1 activity also increased with increasing Gln concentration, but far less than that of TaGS1;2 (Fig. 2C). Only when no N was supplied, TaGS1;1 was found surrounding vessels of xylem (Fig. 5A). These results indicate that TaGS1;1 is also involved in loading Gln to the vascular tissues, but less importance than TaGS1;2. TaGS1;2 activity was inhibited by higher Glu concentration (Fig. 2A), indicating the reverse concentration loading of Gln may be inhibited by high Glu concentration.

The affinity of TaGS1;3 to Glu and hydroxylamine was lower than that of TaGS1;1, but it had the highest V_max_ (Table 1), indicating that TaGS1;3 has strong NH_4_^+^ assimilation ability. The activity of TaGS1;3 was not affected by Gln (Fig. 3C), suggesting that TaGS1;3 did not participate in the reverse concentration loading process of Gln. TaGS1;3 was significantly promoted by the external supply of NH_4_^+^ (Fig. 3C, E, F), and located in pericycle cells of root and leaf mesophyll cells (Fig. 5G, H), indicating that TaGS1;3 mainly performs rapid NH_4_^+^ assimilation at high external NH_4_^+^ concentration, which can prevent the toxicity of high NH_4_^+^ concentration from cells. The root grows rapidly (Table S4) both under medium without N and NO_3_^-^ medium, and a large amount of TaGS1;3 was distributed in the meristematic zone of roots (Fig. 5C, F), indicating that TaGS1;3 was also involved in the assimilation of NH_4_^+^ coming from the metabolism process of rapidly growing roots.

Wheat grows in soils with constantly changing available N forms and concentrations. Therefore, it is difficult for a single TaGS to complete all nitrogen assimilation tasks. Without external nitrogen, the roots no longer absorb inorganic nitrogen, resulting in soluble sugars, carbon skeletons for nitrogen assimilation, accumulation in the roots (Fig. 4B). At the same time, the shoot growth was inhibited due to the lack of nitrogen (Table S4), resulting in the accumulation of photosynthetic products (soluble sugars) in the leaves (Fig. 4B). N stress can cause leaf senescence, promoting proteolysis and N remobilization (Caputo *et al.*, 2009), but roots (meristematic zone) growth was promoted significantly (Table S4) for nitrogen stored in the leaves may be mobilized and translocated through the phloem to the meristematic zone for root growth. During this process, ammonia released by the degradation of nitrogen-containing substances in the leaves was mainly assimilated into Gln by TaGS1;1 located in the mesophyll cells (Fig. 5A), and then loaded into the vessels by TaGS1;1 and TaGS1;2 and distributed around the xylem vessels (Fig. 5A and Fig. 6A). The xylem vessels and phloem sieve tube can exchange substances (Han *et al.*, 1986), so Gln loaded into the vessels can also enter the phloem sieve tube. TaGS1;1 was mainly distributed in vascular bundles (Fig. 5A) to translocate of Gln in the roots. TaGS1;1 and TaGS1;3 were distributed in the meristematic zone of roots (Fig. 5C), and they jointly participated in the assimilation of ammonia produced during the growth and metabolism of cells (Fig. 6A).

**Fig. 6.**
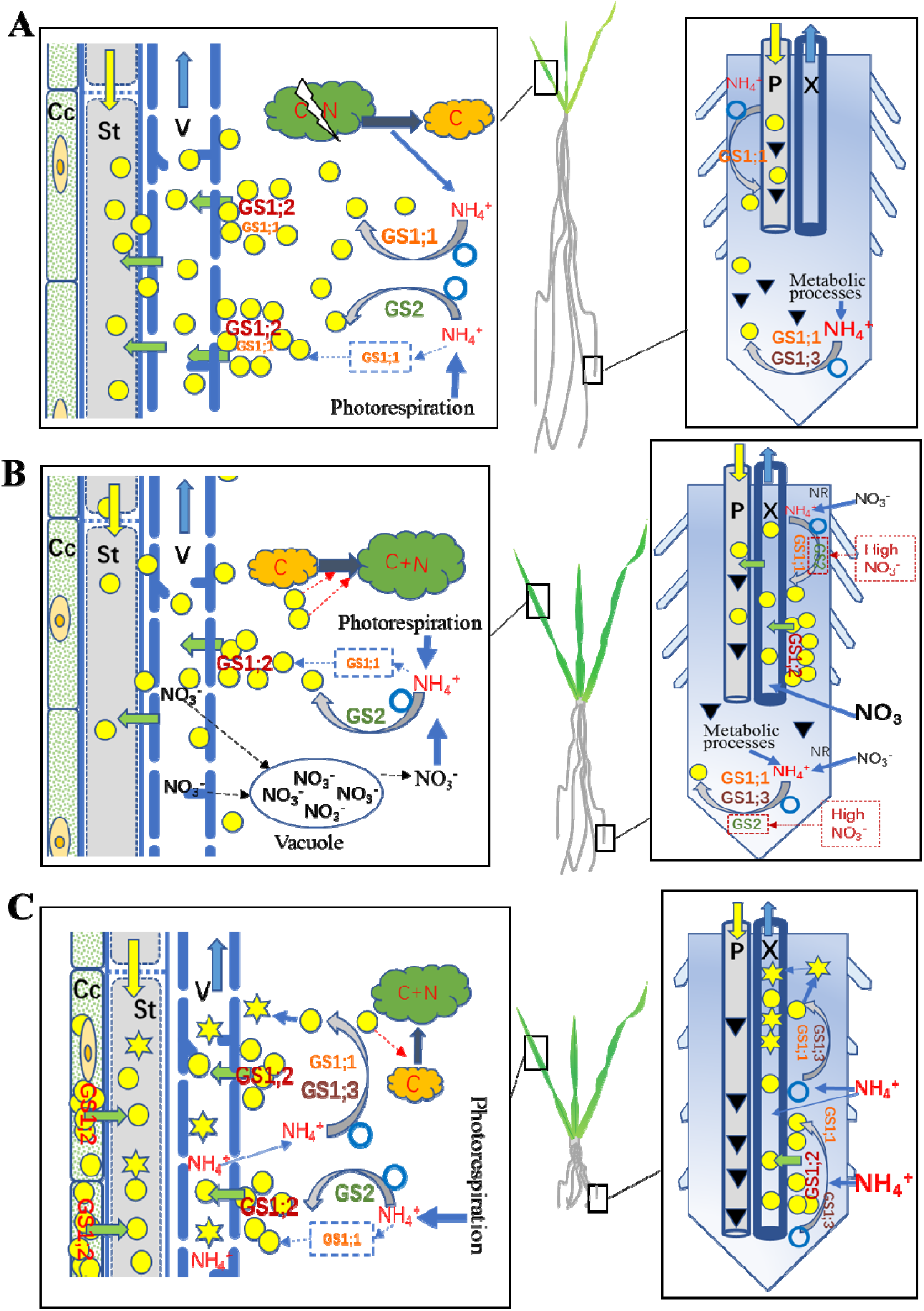
Schematic model of individual TaGS1 synergistically perform nitrogen assimilation and translocation under the condition of N absence (A), NO_3_^-^ supply (B) or NH_4_^+^ supply (C). V, vessel; St, sieve tube; Cc, phloem companion cells; P, phloem; X, xylem.

When NO_3_^-^ was supplied, it will be firstly translocated to the shoot through the xylem and reduced to NH_4_^+^ in the chloroplast of leaves, and then be assimilated into Gln by TaGS2. Some Gln remains in the leaves for leaf growth, some can be loaded into xylem by TaGS1;2 and translocated to other parts through phloem (Fig. 6B). However, the more NO_3_^-^ was supplied, the more TaGS2 was induced in the leaves and roots (Fig. 3D, E, F), and the more NH_4_^+^ was located in the roots (Fig. 4A), indicating that NO_3_^-^ reduction and assimilation occurred both in the leaves and roots. TaGS1;1, TaGS1;3, and TaGS2 were found in the pericycle of maturation zone of roots, whereas TaGS1;2 was mainly distributed around the xylem vessels (Fig. 5E), indicating that Gln was synthesized by TaGS1;1, TaGS1;3, and TaGS2 together, and then translocated to the shoot via xylem by TaGS1;2 (Fig. 6B). Most NO_3_^-^ is translocated to shoot and stored in vacuoles of mesophyll cells or directly stored in vacuoles of root cells (Xu *et al.*, 2012) for a large accumulation of NO_3_^-^ in the shoot and root (Fig. S2). However, the free amino acid content was very low under NO_3_^-^ supply (Fig. 4C) and the content of Gln and Asn (i.e., the organic N translocation forms) were significantly lower than under NH_4_^+^ supply (Fig. 4E, G), indicating that NO_3_^-^ was the most important form for nitrogen translocation and storage.

NH_4_^+^ is an important inorganic nitrogen source, but high NH_4_^+^ concentration tends to produce NH_4_^+^ toxicity to plants (Wang *et al.*, 2016). NH_4_^+^ penetrating into roots has to be immediately assimilated to Gln by GS to prevent NH_4_^+^ toxicity (Funayama *et al.*, 2013). Growing in an environment with a large amount of NH_4_^+^, plants will accumulate a large amount of ammonium (Belastegui-Macadam *et al.*, 2007) and maintain high levels of inorganic nitrogen assimilation in the roots to protect the photosynthetic parts of the plant against ammonium toxicity (Aarnes *et al.*, 2007; Cruz *et al.*, 2006; Hollstein *et al.*, 2010). In our study, with the increase of NH_4_^+^ supply, NH_4_^+^ was accumulated in the root (Fig. 4A) and the root growth was inhibited (Table S4), which allowed the carbon skeleton from the shoot to be used for NH_4_^+^ assimilation. The meristematic zone of roots stopped cell division and differentiated into vascular tissue (Fig. 5I), which helped assimilate products translocation to the shoot in time. During this process, a large amount of TaGS1;1, TaGS1;2, and TaGS1;3 distributed in the root tips consumed the carbon skeleton translocated from the shoot for nitrogen assimilation (Fig. 6C), resulting in a decrease in soluble sugar content in the root (Fig. 4B). A large amount of TaGS1;2 was distributed in the vascular tissue of root tips (Fig. 5I), which helped to load Gln to the vascular tissue (Fig. 6C). Asparagine is another main compound for N storage and translocation due to its high N/C ratio and stability (Ikeda *et al.*, 2004). It is synthesized by asparagine synthase (AS) by the amidation of aspartate (Asp) using Gln as amino donor (Ikeda *et al.*, 2004). As an excellent compound in the carbon economy of nitrogen translocation from roots to shoot, Gln in the root may be transformed into Asn by AS.

When the external NH_4_^+^ exceeds the maximum amount stored and assimilated by the roots, NH_4_^+^ may be translocated to the shoot through the xylem and assimilated in the leaf. In leaves, TaGS1;1 and TaGS1;3 were distributed in the mesophyll cells (Fig. 5G), and they may jointly participate in the assimilation of NH_4_^+^ (Fig. 6). Part of Gln can then be loaded into the vascular tissue by TaGS1;2, distributed in the surrounding vessels of xylem and phloem companion cells (Fig. 5G), and translocated to the tissue short of nitrogen (Fig. 6C).

Based on the above, we can conclude that TaGS1;1 is the primary TaGS1 isozyme for N assimilation, TaGS1;2 mainly participates in the reverse concentration loading of Gln into vascular tissues, TaGS1;3 participates in NH_4_^+^ assimilation of root tip and detoxification of NH_4_^+^, and they are synergistically perform nitrogen assimilation and translocation (Fig. 6). Many studies have suggested that GS1 is closely related to crop nitrogen use efficiency (Funayama *et al.*, 2013; Guan *et al.*, 2015; Martin *et al.*, 2006; Sakurai *et al.*, 1996; Tabuchi *et al.*, 2005; Zhang *et al.*, 2017). However, the outcome of one GS1 overexpression has generally been inconsistent (Thomsen *et al.*, 2014). Considering the synergies of the three TaGS1 isozymes, they should be considered simultaneously to achieve the aim of improving wheat NUE.

## Supplemental date

**Fig. S1** DNA Star multiple alignment of wheat glutamine synthetase amino acid sequences.

**Fig. S2** Effect of NO_3_^-^ supply on the content of NO_3_^-^ in the shoot and root.

**Table S1** The antigenicity, hydrophilic, and surface accessibility of the TaGS isoform specific peptides sequence.

**Table S2** Composition of nutrient solution treated with different nitrogen sources.

**Table S3** List of primers used for qPCR.

**Table S4** Effect of nitrogen regimes on dry weight, fresh weight, root length, and nitrogen content.

## Acknowledgements

We thank the Modern Agricultural Technology System in Henan province (S2010-01-G04) and the 13th five-year national key research and development plan of China (2016YFD0300205 and 2016YFD0300609) for supporting this research.

